# Connections between *Klebsiella pneumoniae* Bloodstream Dynamics and Serotype-Independent Capsule Properties

**DOI:** 10.1101/2025.08.01.668200

**Authors:** Emily L. Kinney, Drew J. Stark, Saroj Khadka, William Bain, Laura A. Mike

## Abstract

*Klebsiella pneumoniae* bacteremia is a significant public health burden with a 26% mortality rate, which increases when the infecting isolate is multi-drug resistant. An important virulence factor of *K. pneumoniae* is the capsule, the protective polysaccharide coat which surrounds the outer membrane and is made up of individual capsular polysaccharide (CPS) chains. Capsule can differ in abundance, attachment, and length of the individual CPS chains. Long, uniform CPS chains are associated with a high level of mucoidy. Typically, mucoid CPS is produced by the hypervirulent *K. pneumoniae* (hvKp) pathotype, which is associated with invasive community-acquired infections. In contrast, the classical *K. pneumoniae* (cKp) pathotype tends to be non-mucoid and is associated with nosocomial infections and multi-drug resistance. There are over 80 serotypes of *K. pneumoniae* capsule. Capsule swap experiments have begun to reveal the effect of serotype on virulence and immune interactions. Clinically, the K2 capsule serotype is a common serotype associated with neonatal sepsis cases. Both cKp and hvKp can produce K2 capsule, but how K2-encoding cKp and hvKp strains differ in a bloodstream infection remains unknown. To fill this gap in knowledge, we characterized the surface properties of K2 serotype cKp and hvKp bloodstream infection isolates, then tested the fitness of these strains in bloodstream infection-related *in vitro* and *in vivo* assays. Understanding how K2 cKp and hvKp strains differ in pathogenic potential provides further insight into how *K. pneumoniae* capsule properties influence bloodstream infection pathogenesis.

**IMPORTANCE:** Many studies have compared the pathogenesis of the two *Klebsiella pneumoniae* pathotypes, but did not control for differences in capsule serotype. In this study, we control for the K2 capsule serotype in classical and hypervirulent pathotypes. Our studies reveal that despite the two pathotypes exhibiting similar human serum survival, they display different tissue tropism in a murine bloodstream infection model. Additionally, although these pathotypes produce similar amounts of capsule, the hypervirulent strains are significantly more mucoid. Our study has provided more evidence on how capsule characteristics impact *K. pneumoniae* bacteremia pathogenesis and the importance of capsule serotype.

## INTRODUCTION

Globally, *Klebsiella pneumoniae* is the third leading cause of antimicrobial resistance-related deaths^3^. It is a Gram-negative pathogen that can infect a variety of host sites, including the urinary tract, lungs, and blood. Bacteremia from *K. pneumoniae* has a 26% mortality rate which increases with antimicrobial resistance^4^. Additionally, according to the Child Health and Mortality Prevention Surveillance network, *K. pneumoniae* is implicated in 1 in 4 deaths among children under age 2 in low-and-middle income countries^5–7^. It is also the primary cause of neonatal sepsis^8–10^. There are two distinctly alarming pathotypes of *K. pneumoniae,* classical (cKp) and hypervirulent (hvKp). cKp strains are generally associated with nosocomial infections and multi-drug resistance^11,12^. Meanwhile, hvKp strains are associated with community-acquired infections and metastatic dissemination^13^. Despite these observed pathogenic differences, both pathotypes can encode similar virulence properties.

Capsule is the protective polysaccharide coat that surrounds the outer membrane of *K. pneumoniae* and is composed of individual capsular polysaccharide (CPS) chains. CPS can differ in serotype (*i.e.* capsule composition), abundance (total capsule synthesized), chain length (mode length of individual CPS chains), chain length diversity (range of CPS chain length), and attachment (outer membrane association). These nuanced capsular properties can contribute to different physiological and pathogenic functions in bacteria^21^. For example, compared to cKp, hvKp has been reported to exhibit increased capsule abundance, mucoidy, and other virulence factors linked with increased frequency of invasive infections^12,22,23^. Recent work demonstrated that capsule abundance is independent of mucoidy, rather more uniform CPS chain length promotes mucoidy^24–26^. While the presence of capsule is a defense mechanism against complement-mediated killing, mucoidy blocks association with macrophages^27–30^.

There are over 80 different *K. pneumoniae* capsule serotypes. However, most of the K-serotype diversity is observed in cKp, as hvKp primarily produces K1 and K2^23^. It should be noted that hvKp isolates have also been found with other capsule serotypes, including K5, K16, K20, K57, and K63^31–36^. In recent years, elegant capsule swap experiments have begun to reveal the effect of these individual serotypes on virulence, immune interactions, and phage dynamics^37–39^. When Huang et al. replaced the K2 capsule biosynthesis locus with either the K1 locus or a complemented K2 locus, the mice exhibited 100% mortality and all intravenous infections had high bacteremia^37^. When the K2 capsule was replaced with the K3 or K23 serotype, the mice exhibited 100% survival and intravenous infections had low bacteremia. Their study highlights the importance of considering capsule serotype in *K. pneumoniae* studies. More broadly, studies comparing cKp and hvKp strains have typically focused on antibiotic resistance profiles, mucoid phenotype, or overall virulence^40–42^. However, most of these comparisons do not control for different capsule serotypes. Since capsule serotype can significantly affect the outcome of an infection, using different serotypes when comparing cKp and hvKp could confound data interpretation^43^. To overcome this limitation, here we compared cKp and hvKp isolates with the same K2 capsule serotype since the K2 capsule serotype is prevalent in neonatal sepsis^8,44^. The K2 serotype is also frequent in both community-acquired and hospital-acquired infections^45^.

Here, we aimed to understand how K2 cKp and hvKp differ in their interaction with host defense mechanisms in the context of bloodstream infections. To assess the role of serotype-independent capsule properties in bacteremia, we studied nine K2 bloodstream isolates (cKp N = 5; hvKp N = 3) and one common K2 hvKp lab strain (ATCC 43816-derived, KPPR1) (**Table 1**). We quantified their capsule properties and their collective fitness in conditions that mimic facets of a bloodstream infection, specifically human serum survival, whole blood survival, host cell association, and dissemination from the bloodstream. Surprisingly, K2 cKp and K2 hvKp exhibited similar phenotypes in all assays, except hvKp strains tend to be more mucoid, associate less with host cells, and achieve higher bacterial burdens in spleens and livers compared to cKp strains. These studies begin to reveal the serotype-independent role of specific capsule properties in a bloodstream infection.

**Table 1.** Summary of K2 isolate genotypic properties. Pathogenwatch was used to predict the listed properties based on whole genome sequencing data^46–49^.

| Isolate | Reference | Pathotype | Sequence Type | *wzi* Type | K Type | *rmp* locus | Siderophores | O Locus | Predicted O Type |
| --- | --- | --- | --- | --- | --- | --- | --- | --- | --- |
| cKp83 | 1 | cKp | 25 | wzi72 | K2 | - | ybt14; ICEKp12 | O1/O2v2 | O1ab |
| KLP203 | 2 | cKp | 25 | wzi72 | K2 | - | ybt14; ICEKp5 | O1/O2v2 | O1ab |
| KLP679 | 2 | cKp | 25 | wzi72 | K2 | - | ybt14; ICEKp5 | O1/O2v2 | O2afg |
| Kp6984 | 14 | cKp | 14 | wzi2 | K2 | - | - | O1/O2v1 | O1ab |
| Kp11996 | 14 | cKp | 14 | wzi2 | K2 | - | - | O1/O2v1 | O1ab |
| KPPR1 | 15-18 | hvKp | 493 | wzi2 | K2 | rmp3 | ybt2; ICEKp1<br>iro3 (truncated) | O1/O2v1 | O1ab |
| KPN165 | 19 | hvKp | 380 | wzi203 | K2 | rmp2 | ybt14; ICEKp12<br>(truncated)<br>iuc2<br>iro2 (truncated) | O1/O2v1 | O1ab |
| hvKp1 | 1,20 | hvKp | 86 | wzi2 | K2 | rmp1 | ybt1-; ICEKp4?<br>iuc1<br>iro1 | O1/O2v1 | O1ab |
| KPN49 | 19 | hvKp | 66 | wzi257 | K2 | rmp2 | ybt12; ICEKp1-iuc2<br>iro2 | O1/O2v2 | O1ab |

## RESULTS

### K2 cKp and hvKp differ in mucoidy, not capsule abundance

hvKp is traditionally described as exhibiting elevated capsule abundance and increased mucoidy when compared to cKp. To test whether this distinction holds while controlling for capsule serotype, we tested the bloodstream infection isolates (**Table 1**) for cell-associated CPS and cell-free extracellular polysaccharide (EPS) production^50^. Unexpectedly, both cKp and hvKp isolates produce similar quantities of CPS and EPS (**Figure 1A-B**). Although mean CPS abundance was the same between the pathotypes, the hvKp strain KPN49 produced more CPS than each cKp strain (**Figure S1A-B**). We then quantified mucoidy using low-speed centrifugation^50^. As expected, the cKp isolates display less mucoidy than hvKp (**Figure 1C**). Notably, we also observed that one cKp strain, Kp11996, was mucoid (**Figure S1A**). Finally, the cell-associated CPS chain length distribution was examined by resolving CPS on an SDS-PAGE gel and visualizing the polysaccharides with Alcian blue followed by a silver stain^50^. Prior work has shown that strains with high mucoidy produce a uniform band of CPS, while non-mucoid strains produce a diverse smear of CPS^24,27,51^. All hvKp strains and the mucoid cKp strain, Kp11996, produce a uniform CPS band, whereas all other cKp strains produce a diverse CPS smear (**Figure 1D**). Notably, the mucoid status of the strain identified by sedimentation assay predicted CPS chain length uniformity by SDS-PAGE, even for cKp strain Kp11996 (**Figure 1C-D**).

**Figure 1.**
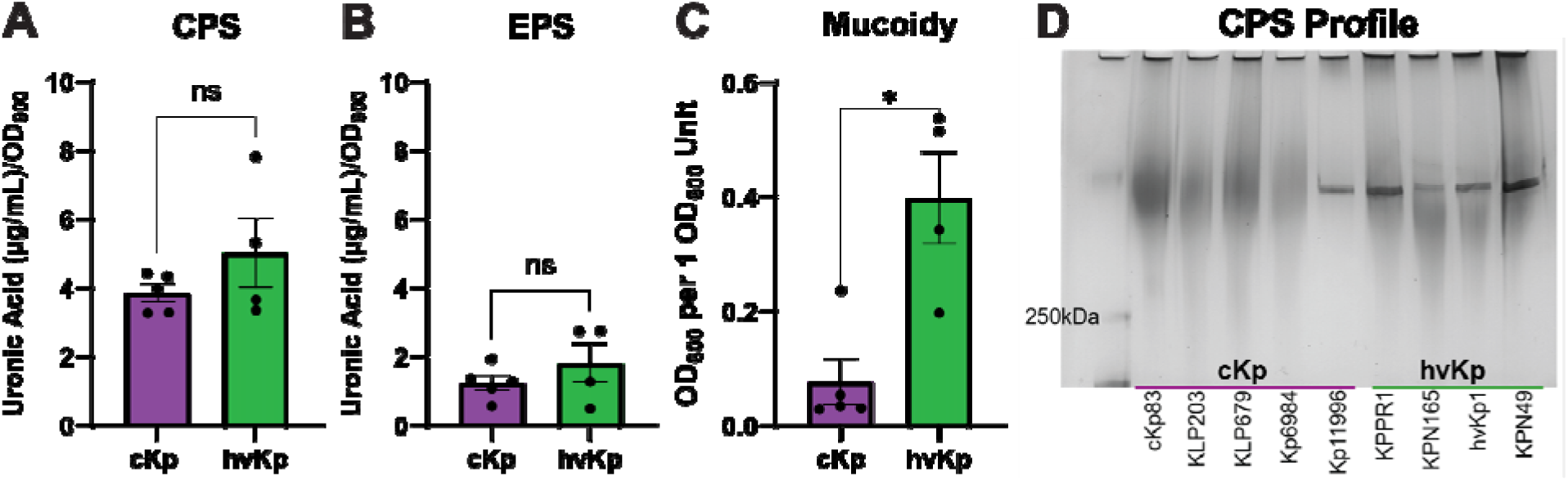
K2 cKp and hvKp strains differ in mucoidy, not capsule abundance. Capsule characteristics of K2 cKp and hvKp strains were determined. Uronic acid quantification measured (**A**) cell-associated CPS and (**B**) cell-free EPS abundance. (**C**) Sedimentation resistance was used to measure mucoidy. (**D**) Purified total CPS was resolved by SDS-PAGE gel to visualize CPS chain length distribution. Shown is a representative image from ≥ 3 independent experiments. (A-C) Each data point represents the average of a single strain, where data were collected in triplicate ≥ 3 independent times. Individual data points for each strain are plotted in Figure S1. Error bars represent the standard error of the mean. To determine statistical significance, a normality test was first applied. A Mann-Whitney test was used for A and an unpaired t-test was used for B and C, where * p < 0.05.

### K2 cKp and hvKp exhibit similar growth kinetics

To determine if growth properties could shape differences in cKp and hvKp pathogenesis, we assessed strain growth in two different media. Strains were cultured in LB or M9 minimal medium with 20% heat-inactivated serum, and the OD_600_ was measured every 15 minutes to generate a growth curve (**Figure S2**). The doubling time and area under the curve were calculated from each growth curve to quantify maximal growth rate and yield (**Figure S3**). No significant differences between cKp and hvKp were detected in either doubling time or area under the curve in either media (**Figure 2**). Therefore, any observed phenotypic differences are not likely due to growth differences.

**Figure 2.**
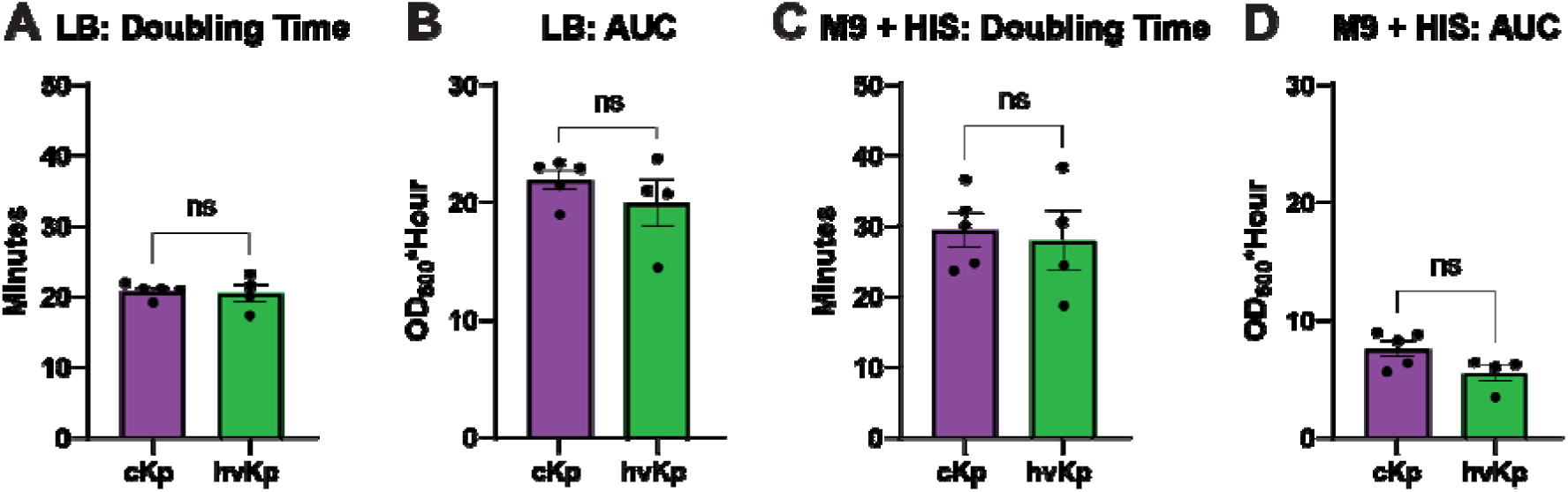
K2 cKp and hvKp exhibit similar growth properties. K2 cKp and hvKp strains were cultured in (**A, B**) LB or (**C, D**) M9 medium with 20% heat-inactivated serum (HIS) as the carbon source. Growth was monitored by OD_600_ for 16 hours. The (**A, C**) doubling time and (**B, D**) area under the curve (AUC) were calculated from each growth curve. Each data point represents the average of a single strain, where data were collected in triplicate ≥ 3 independent times. Individual data points for each strain are plotted in Figure S2 and full growth curves are provided in Figure S3. Bars represent the mean and error bars represent the standard error of the mean. To determine statistical significance, first a test of normality was used. An unpaired t test was used for A, B, and C and a Mann-Whitney test was used for D, ns = not significant.

### K2 cKp and hvKp have comparable human serum resistance

An important part of the immune system in the bloodstream is the complement cascade. In *Streptococcus pneumoniae*, the capsule serotype is a factor in complement resistance^52^. Since our strains are all K2, we assessed whether there was a significant difference between cKp and hvKp sensitivity to pooled human serum when capsule serotype is not a variable. cKp and hvKp survive similarly in human serum, but cKp exhibit a bimodal-like distribution (**Figure 3A**). No killing was observed in heat-inactivated serum, which served as a control to confirm killing of bacteria was due to complement-mediated killing (**Figure 3B**). Finally, a linear regression analysis revealed no significant correlation between serum resistance and any capsule characteristics that we assessed (CPS abundance, EPS abundance, or mucoidy) (**Table 2**). Since the presence of capsule is required for complement-mediated resistance, we predicted that differences in capsule properties might also affect the outcome of this assay, but this was not the case. Furthermore, we predicted that hvKp strains would withstand complement mediated killing better than cKp strains, which could provide one mechanism for their invasive pathogenesis. Instead, when we only consider K2 serotype strains, neither the hvKp pathotype nor CPS abundance predicted serum resistance (**Figure 3** and **Table 2**).

**Figure 3.**
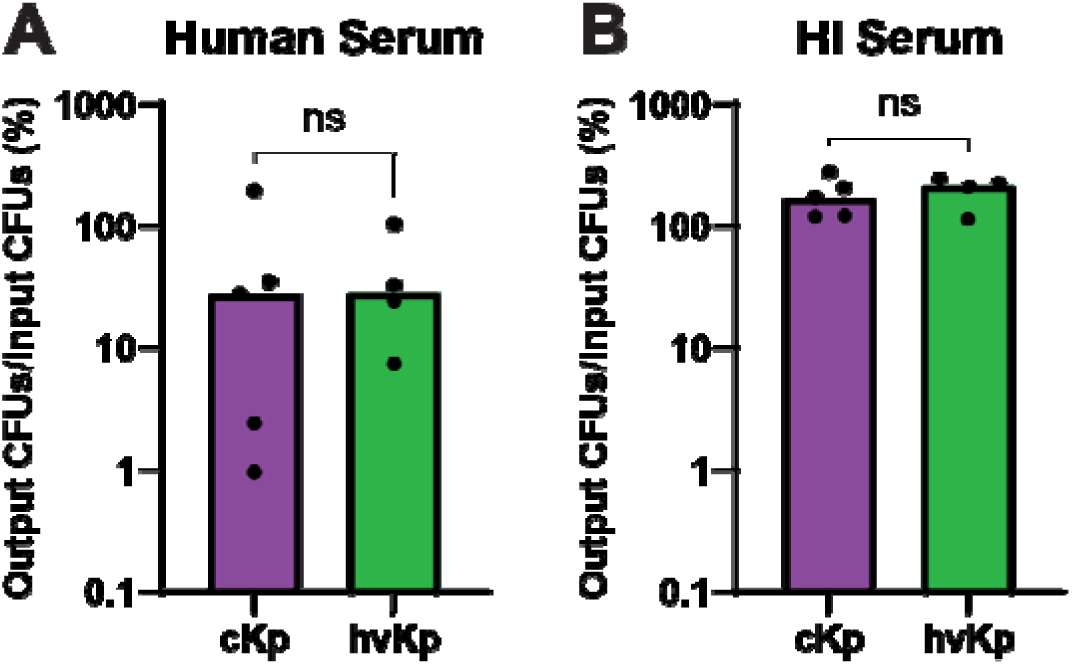
K2 cKp and hvKp have comparable human serum resistance. (**A**) K2 cKp and hvKp were incubated in 90% pooled human serum for 90 minutes. (**B**) Heat-inactivated (HI) serum was used as a control. Data are presented as the percentage of output CFUs divided by the input CFUs. Each data point represents the average of a single strain, where data were collected in triplicate ≥ 3 independent times. Each bar identifies the median. To determine statistical significance, an unpaired t test was used, where ns = not significant.

**Table 2.**
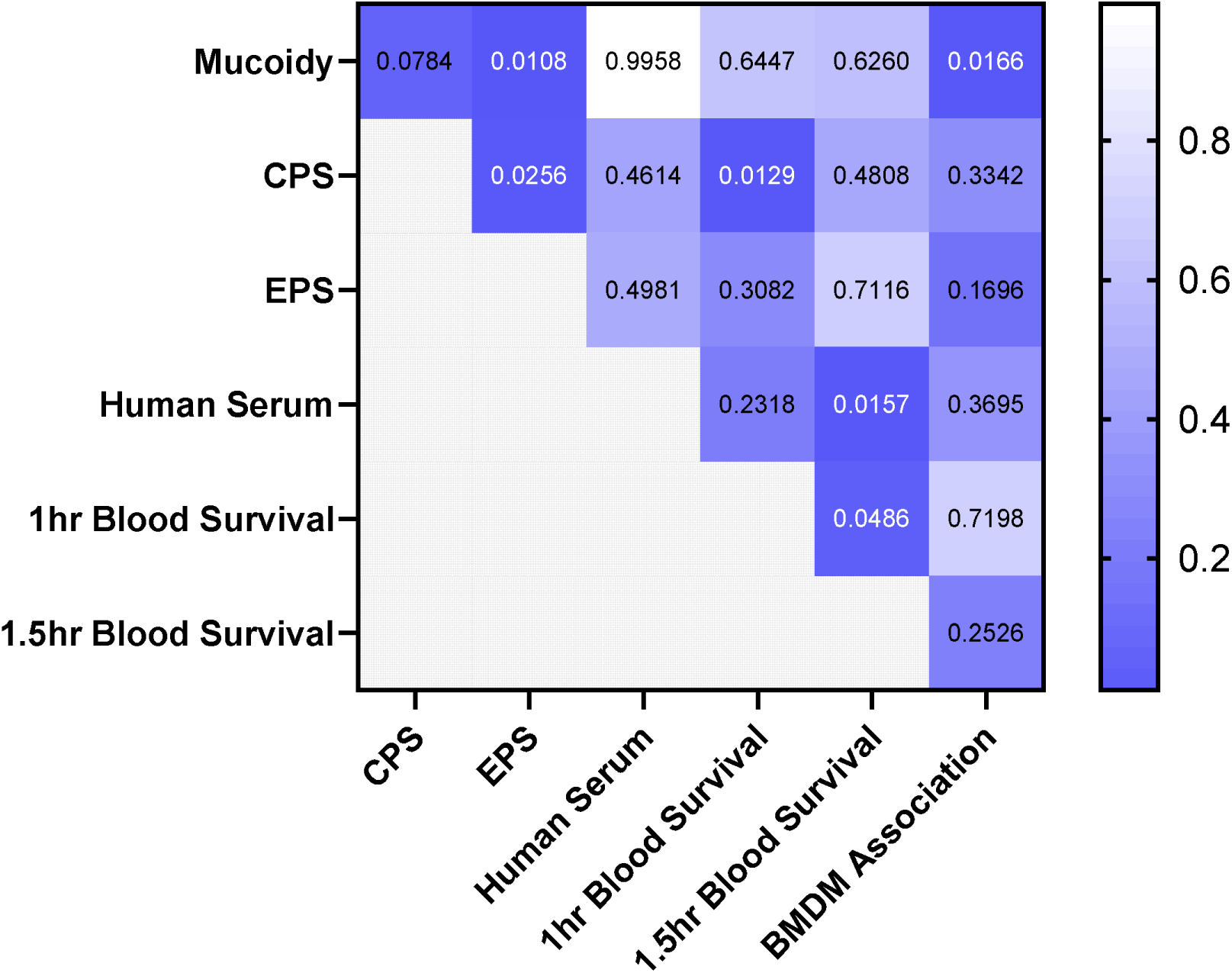
Correlation Between Capsule Properties and Host Interactions. A simple linear regression was performed between every listed assay. The p values are displayed, and p values with numbers in white were deemed significant.

### K2 cKp and hvKp behave similarly in human blood, but cKp associates with macrophages more than hvKp

Given the lack of differences in human serum survival, we next assessed bacterial survival in whole human blood to consider whether another factor in whole blood could influence cKp versus hvKp survival in the bloodstream. cKp and hvKp strains were incubated with human blood for either 1 or 1.5 hours to compare survival patterns of the two pathotypes. cKp and hvKp exhibited similar survival at 1 hour or 1.5 hours (**Figure 4A**). However, there was a non-significant increase in bacteria between the two timepoints, indicating that after an initial period of bacterial death, *K. pneumoniae* can grow in human blood (**Figure 4A**). There was some donor-to-donor variation between strains but no overall patterns (**Figure 5SA-B**). Thus, cKp and hvKp exhibit comparable survival in blood.

**Figure 4.**
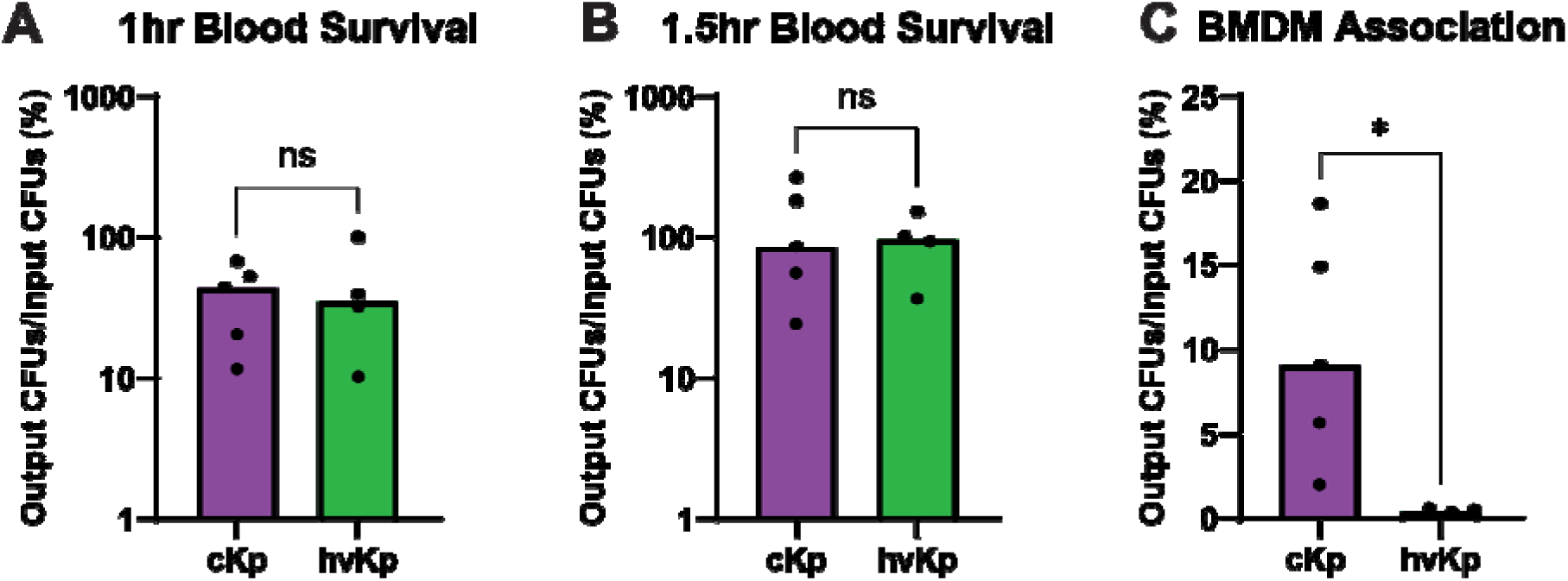
K2 cKp and hvKp survive similarly in human blood, but cKp associates with macrophages more than hvKp. K2 cKp and hvKp were incubated in 90% fresh whole human blood for (**A, B**) 60 or 90 minutes. (**C**) Strains were incubated with immortalized bone marrow-derived macrophages (BMDM) at a MOI of 10 for 2 hours, then washed extensively prior to plating whole BMDM lysates. For all graphs, data are presented as the output CFUs divided by the input CFUs. Each data point represents the average of a single strain, where data were collected in triplicate ≥ 3 independent times. Each bar identifies the median. To determine statistical significance, an unpaired t test (**A,B**) or Mann-Whitney test was used (**C**), where ns = not significant and * p < 0.05.

**Figure 5.**
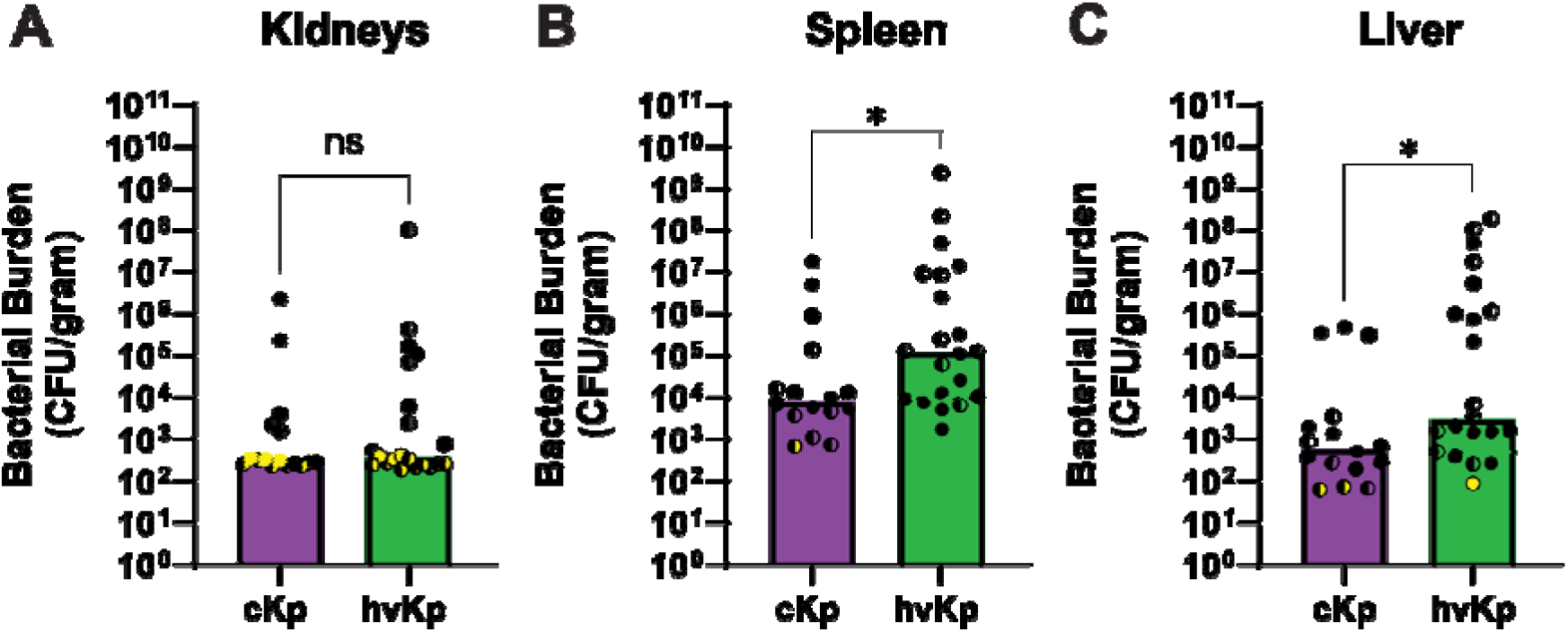
K2 hvKp achieve higher bacterial spleen and liver burdens compared to cKp in a murine bloodstream infection model. C57BL/6 mice were infected with 10^7^ CFU of cKp (Kp6984 or cKp83) or hvKp (KPN165 or hvKp1) strains. After 24 hours, mice were humanely euthanized and bacterial burdens in (**A**) kidneys, (**B**) livers, and (**C**) spleens were enumerated. Bars represent the median and yellow dots represent points below the limit of detection. Each data point represents a single mouse, where half circles identify male mice and full circles identify female mice. Yellow dots represent points below the limit of detection. To determine statistical significance, a Mann-Whitney test was used, where * p < 0.05. Each strain was tested on at least two different days with at least 8 mice.

To determine whether these pathotypes might differ in their interactions with host immune cells, we measured K2 cKp and hvKp association with bone marrow-derived macrophages. Due to antibiotic-resistance profiles, a gentamicin-protection assay could not be performed. Thus, the assay results include both intracellular and cell-bound bacteria. The K2 cKp isolates had significantly higher association (26.3-fold) with the macrophages compared to the K2 hvKp isolates (**Figure 4B**). This difference in susceptibility to macrophage binding suggests that reduced immune cell recognition of hvKp strains could contribute to the increased invasiveness of hvKp strains.

### K2 hvKp achieve higher bacterial spleen and liver burdens compared to cKp in a murine bloodstream infection model

Since K2 strains are prevalent in neonatal sepsis cases and all strains, except KPPR1, were isolated from bloodstream infections, we compared cKp and hvKp isolate dissemination patterns using a murine model of bacteremia^8,44^. Given the more invasive nature of hvKp, we expected to observe greater dissemination of hvKp compared to cKp. Mice were infected with cKp or hvKp isolates through tail vein injection, then blood-filtering organs were harvested after 24 hours. hvKp had significantly higher bacterial burden in the spleen and liver, indicating that the hvKp strains possess increased *in vivo* fitness (**Figure 5A-C**). Additionally, while 100% of cKp-infected mice survived (N = 17/17), only 80.8% of hvKp-infected mice (N = 21/26) survived until the 24-hour timepoint, further supporting the enhanced virulence of hvKp strains *in vivo* (**Figure S6**). The decreased organ burden in cKp-infected mice suggests that cKp are less fit *in vivo*, or alternatively, that hvKp have enhanced capacity for spleen and liver colonization.

## DISCUSSION

The cKp and hvKp pathotypes have been used to categorize *K. pneumoniae* and predict the pathogenic properties of a specific strain. However, even within a single pathotype, the vast genomic variation cannot be understated^53^. This makes it risky to assume phenotypic homogeneity within each pathotype. It has been traditionally thought that hvKp strains are more virulent, have higher capsule abundance, are more mucoid, and disseminate more than cKp. Most studies that have compared these pathotypes have used isolates with a variety of capsule serotypes. We now know that capsule serotype itself is critical for virulence, which confounds these studies because hvKp strains tend to express the more immune-evasive capsule serotypes. The experiments presented here reveal extensive phenotypic overlap between the two pathotypes. Even if a significant difference was detected for a specific phenotype, there was usually at least one strain whose phenotype matched the other pathotype. For example, the mucoidy of cKp strain Kp11996 mirrored the mucoidy of hvKp pathotype strains (**Figure S1A**). The major difference between pathotypes that we observed was the *in vivo* fitness and virulence (**Figure 5 and S6**).

Even though our data indicate that we cannot predict the behavior of individual strains based on pathotype, by controlling for serotype, our relatively small dataset predicts some relationships between surface characteristics and host interactions. Our correlation analyses predict linkages between mucoidy and EPS abundance (*p = 0.0108*), mucoidy and macrophage association (*p = 0.0166*), CPS abundance and EPS abundance (*p = 0.0256*), CPS abundance and early blood survival (*p = 0.0129*), human serum survival and late blood survival (*p = 0.0157*), and early-blood survival and late-blood survival *(p = 0.0486*) (**Table 2**). No other significant correlations between tested phenotypes were detected. It was interesting that EPS abundance had a significant correlation to both CPS abundance and mucoidy. This does not mean that these properties depend on each other; instead, the physical properties are likely the result of related regulatory mechanisms. Since human serum survival correlated with the time-matched blood survival, it seems likely that the same factor drives bacterial killing in human serum and blood.

Compared to cKp, we observed that hvKp strains are more mucoid, associate less with macrophages, and achieve higher spleen and liver burdens, but exhibit no significant differences in CPS or EPS abundance, growth, serum resistance or blood survival (**Figures 1-5**). Combined, these observations suggest that mucoidy is likely to support increased bacterial organ burdens, but more controlled studies are needed due to the amount of genetic variation between the K2 strains examined here. For example, hvKp isolates have additional siderophores that also contribute to *in vivo* fitness (**Table 1**), so we may be indirectly correlating other hvKp factors with behaviors^54–58^. The bloodstream challenges bacterial survival due to phagocyte and complement activity. We used pooled human serum to examine complement resistance and whole human blood to consider other soluble factors in blood that may control bacterial growth. The whole human blood assay captured both bacterial killing (1 hr) and bacterial growth (1.5 hr) (**Figure 4A-B**). This allows us to hypothesize that while blood components can control *K. pneumoniae* growth, some bacteria survive and can use blood as a nutrient source. The two pathotypes did not exhibit differences in human serum or human blood survival (**Figure 3**, **Figure 4A-B**). CPS abundance correlated with early (1 hr), but not late (1.5 hr) survival in blood, suggesting CPS abundance is important for early *ex vivo* blood survival (**Table 2**).

To examine how these two pathotypes differ in regard to phagocyte interactions, we measured the association of cKp and hvKp with immortalized murine bone marrow-derived macrophages (BMDMs). Phagocyte binding of bacteria is an important precursor to phagocytosis. Phagocytosis was not measured here because the standard measure of phagocytosis is a gentamicin-protection assay, and some of the isolates are resistant to gentamicin. A previous study showed that the capsule type controls the ability of liver-resident macrophage Kupffer cells to capture *K. pneumoniae* in the liver (one of the main blood-filtering organs)^37^. Since macrophages are an established line of defense against *K. pneumoniae* infections, we used BMDMs to model bacteria-host cell association. We expect that Kupffer cells or monocyte-derived macrophages would have similar association patterns as the BMDM data reported here. We observed increased BMDM association for cKp compared to hvKp, that inversely correlated with mucoidy, suggesting that mucoidy could shape bacterial binding affinity to host immune cells (**Figure 4C**, **Table 2**). This in agreement with previous literature reporting that when strains are more mucoid, they associate less with macrophages and epithelial cells^25,26,59^.

To compare K2 hvKp and cKp dissemination patterns, we measured dissemination from the blood to the spleen, liver, and kidneys. We picked these organs because of the large volume of blood that filter through them. The hvKp isolates had higher bacterial burdens in the spleen and liver than the cKp isolates (**Figure 5**). The bacterial burdens in these organs do not directly measure primary site dissemination for two reasons. The organ burden does not reflect dissemination from primary sites, but it does reflect bloodstream-based dissemination. Secondly, while increased organ burden could reflect enhanced dissemination from the blood, other factors could contribute to increased *in vivo* survival in that tissue or increased immune evasion^60^. For example, reduced arginine levels in the livers could affect the regulation of hvKp fitness factors^27^. Nonetheless, from the bacterial burden and the overall survival of the mice, we can conclude that the hvKp isolates were more virulent than the cKp isolates tested in our murine model. We cannot conclude whether *in vivo* differences are due to mucoidy, siderophores, another factor, or a combination of these. One interesting observation was the appearance of inflammatory foci on the livers, likely liver abscesses, in both cKp- and hvKp-infected mice. This occurred even though the cKp strains tested did not have a high level of mucoidy or capsule abundance, indicating that another factor may be important for causing liver abscesses. Overall, further studies are needed to determine precisely how mucoidy contributes to bacterial dissemination and association with host cells.

The lack of significant differences between cKp and hvKp regarding serum or blood survival, yet significant differences in BMDM association are potentially insightful. These data may point toward which part of the immune system is important for controlling *K. pneumoniae* bacteremia. While it may be that the capsule levels produced by the strains tested here were all sufficient to protect against complement *in vitro*, another explanation is that human serum resistance may not be critical for limiting *K. pneumoniae* survival in the bloodstream. If this is the case, then dissemination may not occur via the bloodstream. For example, cKp isolate Kp6984, with low mucoidy and low human serum resistance, still has bacteria in every spleen and most of the livers (**Figure S6**). The other cKp isolate tested *in vivo*, cKp83, despite having high human serum resistance, had similar bacterial burden to Kp6984 (**Figure S6**). It was expected that we would see increased human serum resistance from hvKp strains because they are associated with increased dissemination and increased virulence. In order to disseminate to organs other than the initial site of infection, if strains pass through the bloodstream, they must survive. Although complement is a major host factor in the bloodstream, these data indicate it is not a major factor in the pathogenic potentials of cKp and hvKp. Thus, the increased invasiveness of hvKp could be due to differences in immune cell association, not soluble factors. Following this idea, *K. pneumoniae* could hijack macrophages to avoid contact with complement and still disseminate^61,62^.

Our study is limited by the small number of isolates examined. While more isolates may change the outcome of this study, the isolates we have used are from four geographically distinct locations, which makes it more likely for the isolates to represent a range of phenotypes. Although we cannot make firm conclusions between individual strains due to the large number of genetic differences between the strains, these data do identify testable hypotheses for future studies.

A secondary outcome of this study is that we provided further evidence of the importance of capsule serotype in *K. pneumoniae* infections. The overlap of cKp and hvKp strains with the same K2 capsule serotype provides more evidence for the importance of serotype in *K. pneumoniae* pathogenesis. This agrees with capsule swap experiments that have started to uncover the effect of serotype on virulence, immune interactions, and phage dynamics^37–39^. Lee et al. proposed hypotheses on the reason behind why K1 and K2 are more prevalent in hvKp and thus, more virulent^31^. One hypothesis was that K1 and K2 serotypes lack specific mannose-residue repeats recognized by host factors like the mannose-binding receptor on macrophages^38^. Another hypothesis is that K1 has a host-specific monosaccharide sialic acid that allows them to mimic host cells^63^. Studies have already begun to take advantage of this data by strategically making vaccines for important capsule serotypes like K2^64–66^.

In summary, we have revealed that K2 cKp exhibit lower mucoidy, greater macrophage association, and lower bacterial burden after dissemination from the blood compared to K2 hvKp. However, we have also shown that there is a great amount of phenotypic overlap between these two pathotypes. These data offer insights into how K2 capsule characteristics of *K. pneumoniae* impact bacteremia pathogenesis and provide further evidence for the importance of controlling for capsule serotype in *K. pneumoniae* pathogenesis work.

## METHODS

### Strain and Culture Conditions

*Klebsiella pneumoniae* isolates used in this study are reported in **Table 1**. Unless otherwise noted, strains were cultured in low-salt LB medium, or on low-salt LB agar. Liquid cultures were incubated at 37°C for 15.5-16.5 hours at 200 rpm, and plates were incubated at ambient temperatures or 37 °C ^49^.

### Sedimentation Assay

Mucoidy was measured using sedimentation resistance as in Khadka et al^25,50^. The OD_600_ of bacterial cultures was measured and normalized to 1 OD_600_ unit in 1 mL. Then, the samples were subjected to a standard low-speed centrifugation (1,000 *× g* for 5 minutes) and the supernatant OD_600_ was measured. Mucoidy was plotted as the supernatant OD_600_ per 1 OD_600_ unit.

### Uronic Acid Quantification

Capsule abundance was determined by uronic acid quantification as previously described^25,50,67–69^. To start, 250 µL of bacterial culture was added to 50 µL of either 1% Zwittergent 3-14 in 100 mM citric acid (to isolate total CPS) or distilled water (to isolate EPS). CPS samples were incubated at 50°C for 20 minutes. All samples (CPS and EPS) were centrifuged at 17,000 *× g* for 5 minutes, then 100 µL of the supernatant was added to 400 µL ice-cold absolute ethanol. Solutions were incubated on ice for 20 minutes, then centrifuged at 17,000 *× g* for 5 minutes. Pellets were resuspended in 200 µL of water, then incubated at 37°C for 30 minutes. To each sample, 1.2 mL of 0.0125 M sodium tetraborate in sulfuric acid was added, then boiled at 100 °C for 5 minutes and cooled on ice for 5 minutes. The absorbance of acid-treated samples at 520 nm (A_520_) was measured. Subsequently, 10 µL of 0.3% 3-hydroxydiphenyl in 0.125M sodium hydroxide was mixed into each sample and the A_520_ was measured again. A glucuronic acid standard curve was used to determine the uronic acid amount in each sample. EPS was subtracted from total CPS to get cell-associated CPS. CPS and EPS levels were presented as uronic acid concentration per OD_600_.

### CPS Chain Length Visualization

To analyze CPS chain length mode and distribution, purified CPS samples were visualized using an SDS-PAGE gel as previously described^24,50,51,70^. The bacterial cultures were normalized to 1.5 OD_600_. Samples were centrifuged at 21,000 *× g* for 15 minutes, then all the supernatant except for 50 µL (including the pellet) was removed. The remaining pellet was resuspended in 1 mL PBS. The resuspended pellet was centrifuged at 21,000 *× g* for 15 minutes, then all but 250 µL was removed. A 50 µL aliquot of 1% Zwittergent 3-14 in 100 mM citric acid was added to the samples (5:1 sample:Zwittergent), followed by incubation at 50°C for 20 minutes. CPS samples were pelleted at 17,000 *× g* for 5 minutes, then 100 µL of the supernatant was added to 400 µL ice-cold absolute ethanol. Solutions were incubated on ice for 20 minutes, then pelleted at 17,000 *× g* for 5 minutes. Pellets were rehydrated in 200 µL of water at 37°C for 30 minutes. Next, 75 µL of solubilized CPS samples were added to 25 µL of 4x SDS-gel loading dye, then 20 µL of each sample was resolved on a 4-15% Mini-PROTEAN TGX stain-free pre-cast gel (Bio-Rad) by applying 300 V for 4.5 hours at 4°C. The gel was washed five times in ultra-pure water for 10 minutes, then stained with 0.1% Alcian blue in stain base solution (40% ethanol and 60% 20 mM sodium acetate, pH 4.75) for 1 hour, then stain base solution was added to de-stain the gel overnight. The gel was stained using a Pierce Silver Stain Kit (ThermoFisher) and imaged on a Bio-Rad GelDoc.

### Bioinformatics

Pathogenwatch software (version 23.4.0 https://pathogen.watch) was used to determine the sequence type, O locus, and O type from the genomic sequence of the bacterial strains^46–49^. The genomic sequences were obtained from either the NCBI public database, requested from the strain source, or sequenced on the Illumina platform (SeqCoast).

### Bacterial Growth Assay

Bacterial growth curves were generated following the protocol detailed by Pariseau et al^49^. Specifically, stationary phase bacterial cultures were sub-cultured in LB and incubated for 1.5 hours at 37 °C shaking at 200 rpm. The sub-cultured samples were diluted to 0.0001 OD_600_ in either LB or low-iron M9 minimal medium supplemented with CaCl_2_, MgSO_4_, and 20% heat-inactivated human serum (Innovative Research). The samples were cultured with continuous shaking at 37 °C and the OD_600_ was measured every 15 minutes for 16 hours using a microplate reader (Biotek Synergy HTX by Agilent).

### Human Serum Survival

Bacterial survival in human serum was measured as described by Mike et al^25^. Strains were centrifuged at 16,249 *× g* for 10 minutes then resuspended in sterile PBS. The resuspended samples were then diluted to a OD_600_ of 0.02. The diluted bacterial preparations were incubated in 90% pooled human serum (Innovative Research) or heat-inactivated serum (inactivated at 58 °C for 1 hour) at 37 °C for 90 minutes. Bacterial colony forming units (CFUs) in the input and output samples were enumerated by serial dilution and plated on LB agar. Bacterial survival was presented as the percentage of input CFUs surviving after incubation.

### Human Blood Collection

Blood from healthy human donors was collected to use in the whole blood survival assay and the isolation of human monocytes. Approximately 30 mL of blood were obtained through venipuncture after informed consent of the donor (CR19040346-008). The blood samples were collected in a sterile collection tube containing ACD-A as an anti-coagulant.

### Survival in Whole Blood

Overnight bacterial cultures (1 mL) were centrifuged at 21,000 *× g* for 15 minutes then resuspended in 1 mL sterile PBS, followed by dilution to a OD_600_ of 0.02. Then, 10 µL of bacterial sample was incubated in 90 µL human blood (ACD-A tube) statically at 37°C for 60 or 90 minutes. Bacterial CFUs in the input and output samples were enumerated by serial dilution and plated on LB agar. Bacterial survival and/or growth were presented as the percentage of input CFUs present after incubation.

### Bone Marrow-Derived Macrophage Association Assay

Bone marrow-derived macrophages (BEI Resources NR-9456) were used to measure bacterial association to host cells^24^. The macrophages were grown in a 24-well plate (*n* = 200,000 cells per well). Then, each well was infected with bacterial cells prepared in 1 mL DMEM medium at an MOI of 10.The infected macrophages were briefly centrifuged down (54 x *g* for 5 minutes) to initiate cell contact. Then, the plate was incubated for 2 hours, followed by washing with 1 mL sterile PBS three times. After washing, 1 mL 0.2% Triton-X-100 was added to each well for 5 minutes while shaking to lyse monocytes for bacterial release. The inocula (input) and monocyte lysates (output) were serial diluted and CFUs enumerated on LB agar. Bacterial association was presented as the percentage of output CFUs relative to input CFUs.

### Murine Bloodstream Infection Model

The *K. pneumoniae* murine bloodstream infection model used in this study was adapted from a previously described model and performed in adherence to humane animal handling recommendations and approved by the University of Pittsburgh Institutional Animal Care and Use Committee (protocol: 24105753)^71^. C57Bl/6 mice (7-8 weeks old) were procured from Charles River Laboratory (Ashland, OH, USA). The mice were injected with 10^7^ bacteria via tail vein. After 24 hours, the mice were humanely euthanized. Liver, spleen, and kidneys were collected then homogenized in sterile PBS. Bacterial burdens were determined by serial diluting the homogenized organs, then enumerating CFUs on LB agar. Bacterial burden was presented as the log transformed CFUs divided by organ weight.

### Statistics

All data, except for the murine bloodstream model, were collected on at least three different days. For the murine model, bacterial strains were used at least two different days. All statistical analyses were computed in Prism 10.3.1 (GraphPad Software, La Jolla, CA, USA). Tests of normality were performed on all data. If normal, an unpaired t test was used to compare data. If not normal, a Mann-Whitney test was used. For data with more than two comparisons, a one-way ANOVA was used if the data were normal, and a Kruskal-Wallis test was used if the data were not normal. Male and female mice and human blood donors were used to consider sex as a variable.

## ACKNOWLEDGEMENTS

The following reagent was obtained through BEI Resources, NIAID, NIH: Macrophage Cell Line Derived from Wild-Type Mice, NR-9456. We thank Drs. Tom Russo, Mike Bachman, Travis Kochan, Alan Hauser, Lora Pless, and Lee Harrison for the strains in this publication. We thank members of the Bain lab for their help with blood collection. We thank members of the Mike lab and Program in Microbiology and Immunology at the University of Pittsburgh for critical feedback, especially Brooke Ryan, Grace Shepard, Drs. Daria Van Tyne, John Alcorn, Prabir Ray, and Tera Levin.

Research reported in this publication was supported by the University of Pittsburgh College of Medicine and American Heart Association 23CDA1056712 (L.A.M.). This content is solely the responsibility of the authors and does not necessarily represent the official views of the American Heart Association.

